# Maturation of Nucleus Accumbens Synaptic Transmission Signals a Critical Period for the Rescue of Social Deficits in a Mouse Model of Autism Spectrum Disorder

**DOI:** 10.1101/2023.02.08.527742

**Authors:** Melina Matthiesen, Abdessattar Khlaifia, Carl Frank David Steininger, Maryam Dadabhoy, Unza Mumtaz, Maithe Arruda-Carvalho

## Abstract

Social behavior emerges early in development, a time marked by the onset of neurodevelopmental disorders featuring social deficits, including autism spectrum disorder (ASD). Although deficits in social interaction and communication are at the core of the clinical diagnosis of ASD, very little is known about their neural correlates at the time of clinical onset of the disorder. The nucleus accumbens (NAc), a brain region extensively implicated in social behavior, undergoes synaptic, cellular and molecular alterations in early life, and is particularly affected in ASD mouse models. To explore a link between the maturation of the NAc and neurodevelopmental deficits in social behavior, we compared age-dependent changes in spontaneous synaptic transmission in NAc shell medium spiny neurons (MSNs) between the highly social C57BL/6J mouse strain and the idiopathic ASD mouse model BTBR *T^+^ Itpr3^tf^/J* at postnatal day (P) 4, P6, P8, P12, P15, P21 and P30. We found that MSNs from both C57BL/6J and BTBR mice display age-dependent increases in spontaneous excitatory and inhibitory synaptic currents between P4 and P30. Comparison of NAc spontaneous transmission between strains showed that BTBR MSNs display increased excitatory transmission during the first postnatal week, and increased inhibition across the first, second and fourth postnatal weeks, suggesting accelerated maturation of excitatory and inhibitory synaptic inputs onto BTBR MSNs compared to C57BL/6J mice. These early life changes in synaptic transmission are consistent with a potential critical period in the maturation of the NAc, which could maximize the efficacy of interventions affecting social behavior. To test this possibility, we treated BTBR mice in either early life (P4-P8) or adulthood (P60-P64) with the mTORC1 antagonist rapamycin, a well-established rescue intervention for ASD-like behavior. We found that rapamycin treatment rescued social interaction deficits in BTBR mice when injected in infancy, but not in adulthood. These data emphasize the importance of studying brain regions involved in the pathophysiology of neurodevelopmental disorders at clinically-relevant time points, which may offer novel insight into the timing and targets of therapeutic interventions to maximize positive outcomes.

## Introduction

Social behaviors, which include social communication, interaction, parental and reproductive behaviors^1,2^, enable intraspecies interactions necessary to find food, mate and avoid predation^3^. Consistent with their key role in survival, early deficits in the development of social interaction and communication have devastating consequences^45^, and are a hallmark of several neurodevelopmental disorders^6^, including Fragile X syndrome^7^, schizophrenia^8^, and autism spectrum disorder (ASD)^9^. ASD encompasses a set of heterogeneous neurodevelopmental disorders affecting an estimated 1% of the world population^9^, and features impairments in social cognition, communication and social perception^10^. Even though deficits in social interaction and communication are at the core of clinical diagnoses for neurodevelopmental disorders such as ASD^11^, very little is known about the neural basis of such deficits, particularly at the time of clinical onset and diagnosis^12^.

Evidence points to conserved brain regions underlying social processing across species^13^. In particular, the rodent nucleus accumbens (NAc) and its projections^14,15^ have been extensively implicated in multiple facets of social behavior, including social interaction and motivation^16–18^, social cognition^19^, social approach^20,21^, social memory^22–24^, social-spatial representations^25^ and susceptibility to social stress^26–30^. Consistent with its important role in social behavior, imaging studies of ASD patients show abnormal activity in NAc^31,32^, as well as changes in functional connectivity within frontostriatal^33–35^ and striatal^36^circuits. Accordingly, ASD mouse models display altered NAc synaptic plasticity and transmission^37–39^, as well as changes in gene expression of inhibitory and excitatory markers^40^ in adult animals. Importantly, manipulations targeted to the NAc can mimic^41^ or rescue behavior deficits in a range of ASD mouse models^37,42–44^, further implicating the NAc in ASD pathophysiology.

Recent evidence has highlighted the value and long-term benefits of early ASD interventions^45,46^, but the precise time window of optimal efficacy for these interventions has not been clearly defined. In the developing sensory cortex, plasticity in response to external stimuli is curtailed within critical periods^47^, suggesting that similar mechanisms could guide the efficacy of rescue interventions in other brain regions. As with sensory cortex development, the rodent NAc also undergoes early postnatal changes in synaptic transmission^48–52^ and plasticity^53^, in the levels of dopamine receptors^54–57, 58^, dopamine/serotonin transporters^59,60^, stimulus-evoked dopamine transients^61^, complement C3^58^ and microglia^62^, which stabilize around the start of adolescence. To determine whether the timing of these developmental changes defines a window of increased efficacy for interventions, they must be examined in animal models featuring early social deficits, paralleling the clinical presentation of ASD.

Several studies have identified rapamycin, an inhibitor of the serine/threonine kinase mammalian target of rapamycin (mTOR), as an effective pharmacological intervention in ASD animal models^63–66^. mTOR is a critical regulator of cell growth, proliferation, and metabolism^67,68^, as well as synaptic plasticity and memory consolidation^69–71^. mTOR is part of two structurally and functionally distinct multiprotein complexes called mTOR complex 1 (mTORC1) and mTOR complex 2 (mTORC2), of which only mTORC1 shows sensitivity to acute inhibition by rapamycin^68^. Importantly, impaired mTORC1 signaling has been implicated in ASD and related neurodevelopmental disorders^67,72–74^, consistent with the success of rapamycin treatment in rescuing social behavior deficits in ASD mouse models^75–77^.

Here, we compared the maturation of spontaneous excitatory and inhibitory synaptic transmission of NAc shell medium spiny neurons (MSNs) of C57BL/6J mice and a mouse model of idiopathic ASD, the BTBR T^+^*Itpr3^tf^/J* (BTBR)^78,79^ mouse strain. BTBR mice display face validity to all three core ASD behavioral symptoms: deficits in social communication^80–82^, deficits in social interaction^79,83–85^ and repetitive behaviors^79,84–86^. Specifically, when compared to the highly social C57BL/6J strain, BTBR mice display changes in ultrasonic vocalizations, a well-established metric of social communication^87,88^, starting from infancy, with a remarkably reduced repertoire of calls as early as postnatal day (P)8^80^, matching the early social deficits seen in ASD patients^6^. We found that both strains show age-dependent increases in NAc shell MSN spontaneous excitatory and inhibitory transmission, particularly at the start of the second postnatal week. Compared to C57BL/6J, BTBR mice show changes in spontaneous transmission from as early as P4, with increased excitation during the first postnatal week and increased inhibition in the first, second and fourth weeks. Based on these data, we next tested the efficacy of targeting treatment with the mTORC1 inhibitor rapamycin to the age of onset of maturational changes in NAc synaptic transmission. We found that rapamycin treatment was successful at rescuing BTBR deficits in social interactions when applied in infancy, but not in adulthood, suggesting that early changes in synaptic transmission may act as a marker for a critical period maximizing the efficacy of therapeutic interventions.

## Materials and Methods

### Animals

C57BL/6J (Jackson Laboratory) and BTBR *T+ Itpr3tf/J* (Jackson Laboratory; referred to as BTBR for simplicity) mouse strains were bred at the University of Toronto Scarborough and kept on a 12h light/dark cycle (lights on at 07:00 h) with access to food and water *ad libitum*. Date of birth was assigned P0. Approximately equal numbers of females and males were used for social behavior, and males were used for slice electrophysiology experiments. Offspring were weaned with same-sex siblings on P21 (2-4 mice per cage). All animal procedures were approved by the Animal Care Committee at the University of Toronto.

### Stereotaxic Surgery

C57BL/6J and BTBR male mice underwent surgery at P8 or P23. Mice were given an intraperitoneal (IP) injection of ketamine (100 mg/kg) and xylazine (5 mg/kg) and placed in a stereotaxic frame (Stoelting). AAV1.CaMKIIa.hChR2(H134R)-eYFP.WPRE.hGH virus (Addgene) was infused bilaterally in the infralimbic cortex (IL; coordinates relative to bregma: P8: AP +1.2 mm, ML ± 0.25 mm, and DV −1.7 mm; P23: AP +1.8 mm, ML ± 0.25 mm, and DV −2.6 mm). Viral particles (0.2 μl/hemisphere) were delivered at a rate of 100 nl/minute by way of a motorized microsyringe. After viral infusion, the needle was left in place for five additional minutes to allow for diffusion of the virus, and then was slowly removed from the injection site. Mice were then returned to their home cages for seven days to allow for recovery and viral expression. Whole-cell patch-clamp recording experiments were performed at P15 or P30.

### Slice Electrophysiology

Electrophysiological recordings were obtained from male C57BL/6J and BTBR mice at different postnatal ages (P4, P6, P8, P12, P15, P21 and P30). Mice were deeply anesthetized using isoflurane and the brain was quickly removed and placed in ice-cold sucrose-based cutting solution containing (in mM): 180 sucrose, 2.5 KCl, 1.25 NaH_2_PO_4_, 25 NaHCO_3_, 1 CaCl_2_, 2 MgCl_2_, 2 Na^+^ pyruvate and 0.4 L-Ascorbic acid with 95% O2 and 5% CO_2_. Coronal brain slices containing the NAc (250 μm-thick) were prepared with a vibratome (Leica VT1000S). Slices were then incubated for 30 min at 30 °C in a recovery solution composed of 50% sucrose-based cutting solution and 50% artificial cerebrospinal fluid (ACSF). The ACSF was composed of (in mM): 120 NaCl, 2.5 KCl, 1.25 NaH_2_PO_4_, 25 NaHCO_3_, 11 glucose, 2 CaCl_2_, 1 MgCl_2_, 2 Na^+^ pyruvate and 0.4 L-Ascorbic acid with 95% O_2_ and 5% CO_2_. Slices were then placed in regular ASCF for another 30 min at room temperature before electrophysiology recordings. After recovery, slices were placed in the recording chamber and perfused with 2ml/min oxygenated ACSF at room temperature for recordings.

Whole-cell patch-clamp recordings were obtained from NAc shell MSNs using borosilicate glass pipettes (3-5 ΩM; WPI) filled with intracellular solution containing (in mM): 120 Cs-methanesulfonate, 10 HEPES, 0.5 EGTA, 8 NaCl, 4 Mg-ATP, 1 QX-314, 10 Na-phosphocreatine, and 0.4 Na-GTP, pH 7.3, at 290 mOsmol. Spontaneous excitatory (sEPSCs) and inhibitory (sIPSCs) postsynaptic currents were recorded from the same cell at holding potentials of −60 mV and 0 mV, respectively, and five minutes of stable recordings were analyzed for frequency and amplitude (miniAnalysis; Synaptosoft).

For *ex vivo* optogenetic experiments, IL axon terminals were stimulated using TTL-pulsed microscope objective-coupled LEDs (460 nm, ~1 mW/mm2; Prizmatix) and light-evoked EPSCs were recorded from MSNs at holding potential of −60 mV. Glutamate release probability from IL terminals to NAc shell MSNs was assessed by measuring paired-pulse ratio by dividing the amplitude of the second light-evoked EPSC by the amplitude of the first EPSC at inter-stimulus intervals of 250 and 100 ms for P15 and P30 respectively. All electrophysiology recordings were obtained in the absence of synaptic blockers.

Electrophysiology data were acquired using a MultiClamp 700B (Molecular Devices), digitized at 10 kHz using Digidata 1550A and pClamp 10 (Molecular Devices). Recordings were low pass filtered at 4 kHz. Access resistance was regularly assessed during experiments and data were included only if the holding current was stable and access resistance varied <20% of initial value.

### Behavior

The three-chamber social interaction and social memory tests were conducted following established protocols from the Crowley group^89^. Tests took place in a 60×20×40cm plexiglass apparatus with three 20×20×40cm chambers interconnected by two 7×40cm vertically removable plexiglass doors. Two 8.5×8.5×40 cm enclosures with three 0.3cm vertical slits per side were placed at the lateral wall of the right and left chambers and used to hold social target animals as required in the tasks. The chamber was elevated 41 cm off the floor and a camera was mounted 75 cm above the chamber on a metal rack.

On the day prior to behavior testing, C57BL/6J and BTBR mice were socially isolated for an hour in a separate room. On the day of testing, mice were socially isolated for an hour, followed immediately by habituation to the apparatus for 10 minutes. Following habituation, mice underwent the three-chamber social interaction test, in which an age-, strain- and sex-matched stranger mouse was placed in one of the enclosures in a counterbalanced manner. Experimental mice were placed in the middle chamber at the start of the experiment, and opening of the doors marked the start of the social interaction test. Mice were able to freely explore the three chambers for a total of 10 minutes.

Social memory tests took place one hour after the social interaction test. For the social memory tests, two mice were placed in the enclosures: the stranger mouse used in the social interaction test (now familiar mouse), and a novel age, strain and sex-matched mouse, that the experimental mouse had never previously encountered (stranger mouse). The experimental mouse was placed in the middle chamber with the doors closed. Following the opening of the doors, the animal was free to explore the three chambers for 10 minutes.

Social behavior was conducted during the late phase of the light cycle. A subset of animals had undergone social interaction and social memory tests in the dark cycle, but similar to what has been described in the literature^90^, we saw no behavioral differences between light and dark cycles for these strains, and therefore consolidated the datasets. Behavior was analyzed using ANY-maze^®^ software, and cross validated through manual scoring by an experimenter blind to experimental conditions.

### Drugs

The stock solution of rapamycin (50 mg/kg in 100% ethanol; LC laboratories; Woburn, MA; U.S.A) was stored at −80°C. Stock solution was diluted in 10% of polyethylene glycol 400 (PEG400) and 10% Tween 80 to a final concentration of 1mg/ml in 2% ethanol and stored at −20°C ^66^. Mice received an intraperitoneal injection of rapamycin (0.5mg/kg) or vehicle (10% PEG400, 10% Tween; 1mg/kg)^66^ for five consecutive days, from either P4 to P8 or P60 to P64.

### Perfusions and Sectioning

BTBR mice were treated with vehicle or rapamycin from P4-P8 and were perfused at P30 for immunohistochemistry against phosphoS6. At P30, behaviorally naïve mice were injected with Avertin (250mg/kg, i.p.) and once deeply anesthetized, were transcardially perfused with 0.1M phosphate buffered saline (PBS), followed by 4 % paraformaldehyde (PFA). The brains were then extracted and stored overnight at 4°C in 4% PFA. The brains were sectioned using a vibratome (VT 1000, Leica) to obtain 50 μm coronal sections, which were stored in a 60% glycerol and 0.01% sodium azide in PBS solution at −20°C.

### Immunohistochemistry

Sections containing the NAc shell were treated with 0.3% Triton X-100 in PBS for 15 minutes, followed by 10% normal goat serum in 0.1% Triton X-100 in PBS for 1 hour. Sections were incubated with rabbit monoclonal anti-phospho-S6S^240/244^ (pS6; 1:1000; Cell Signaling, Beverly, MA, RRID:AB 10694233) for 48h, followed by secondary antibody incubation with Alexa Fluor 594-conjugated goat anti-rabbit IgG (1:500; Jackson Immunoresearch Laboratories) and Hoechst dye (1:1000; Thermo Fisher Scientific) for 90 minutes. Sections were mounted on glass slides (Thermo Fisher Scientific) with Permafluor aqueous mounting medium (Thermo Fisher Scientific) and coverslipped for storage at 4°C.

### Image Acquisition and Quantification

Image acquisition was performed using a Nikon Eclipse Ni-U epiluorescence microscope. Immunofluorescence was visualized with an LED illumination system (X-Cite 120 LED Boost, Excelitas Technologies) and captured with a Nikon DS-Qi2 digital camera. All images were acquired at 10x magnification using Plan-Apochromat differential interference contrast (DIC) N1. When brightness and/or contrast adjustments were made in a figure, these changes were made equally to all photomicrographs.

Cell counts were done manually using Fiji/ImageJ software (v. 1.53e) by an experimenter blind to treatment conditions. Three to five sections per subject containing the NAc shell were used for cell counts. For each section, the integrated density in the NAc shell was acquired, corrected for background, and averaged to obtain the mean integrated density per animal.

### Statistical Analysis

Data are presented as mean ± standard error of the mean. All statistical analyses were performed in Graphpad Prism^®^ version 9. Potential sex differences were first assessed using a two- or three-way, repeated measures ANOVA and given the absence of effects, data were pooled for subsequent analyses. Exploration time in all chambers in the social interaction and social memory tests was analyzed by two-way, repeated-measures ANOVA followed by Sidak’s post-hoc tests. Distance travelled was analyzed using unpaired two-tailed t tests. For electrophysiology experiments and immunohistochemistry, paired t-tests and Wilcoxon signed-rank tests were used for within-group comparisons, unpaired t-tests and one- or two-way repeated measures ANOVA followed by Tukey’s and Sidak’s post-hoc tests respectively were used for normally distributed data and Mann-Whitney Rank Sum test when data did not pass normality. For all analyses, p<0.05 was considered significant.

## Results

### Maturation of spontaneous excitatory currents of medium spiny neurons of the nucleus accumbens shell of C57BL/6J and BTBR mice

To examine the maturation of spontaneous excitatory and inhibitory synaptic transmission of the NAc shell in C57BL/6J and BTBR mice, whole cell patch-clamp recordings were obtained from visually identified MSNs (Fig. 1A). In MSNs from both strains, spontaneous excitatory postsynaptic currents (sEPSCs) were observed as early as P4 and developed gradually during the first postnatal month (Fig. 1). In C57BL/6J mice, sEPSC frequency showed a significant increase at P15 and P21 with a further increase at P30 (Fig. 1B-C, one way ANOVA, F_(6, 64)_ = 25.40 p<0.0001, Tukey’s multiple comparison tests: P4 vs P15, p=0.02; P4 vs P21, p=0.0003; P4 vs P30, p<0.0001; P6 vs P15, p=0.04; P6 vs P21, p=0.0004; P6 vs P30, p<0.0001; P8 vs P21, p=0.0017; P8 vs P30, p<0.0001; P12 vs P30, p<0.0001; P15 vs P30, p<0.0001; P21 vs P30, p<0.0001). However, in BTBR mice, a marked increase in sEPSC frequency was observed at P12, with a further increase from P21 (Fig. 1B, D, one way ANOVA, F_(6, 66)_ = 14.86 p<0.0001, Tukey’s multiple comparison tests: P4 vs P12, p=0.007; P4 vs 21 and P30, p<0.0001; P6 vs P21, p=0.0001; P6 vs P30, p<0.0001; P8 vs P21, p=0.0007; P8 vs P30, p<0.0001; P12 vs P30, p=0.0007; P15 vs P21, p=0.004; P15 vs P30, p<0.0001). Interestingly, sEPSC amplitude displayed a distinct maturation profile. While C57BL/6J mice showed increased sEPSC amplitude between P6-P8 and P15 and P30 (Fig. 1E, one way ANOVA, F_(6, 66)_ = 7.429 p<0.0001, Tukey’s multiple comparison tests: P6 vs P15, p=0.0004; P6 vs P30, p=0.0005; P8 vs P15, p=0.0003; P8 vs P30, p=0.0003), the amplitude of sEPSCs in MSNs from BTBR mice was decreased at P8, P12 and P21 relative to P4 (Fig. 1F, one way ANOVA, F_(6, 70)_ = 3.480 p=0.0046, Tukey’s multiple comparison tests: P4 vs P8, p=0.002; P4 vs P12, p=0.03; P4 vs P21, p=0.01). Furthermore, sEPSC time constant displayed a marked increase during the second (P12, P15) and third (P21) postnatal weeks in C57BL/6J mice and similarly during the second (P8) and third (P21) postnatal weeks in BTBR mice (Fig. 1G, one way ANOVA, F_(6, 67)_ = 5.602 p<0.0001, P4 vs P12, p=0.006; P4 vs P15, p=0.0006; P4 vs P21, p<0.0001; Fig. 1H, one way ANOVA, F_(6, 70)_ = 3.121 p=0.0091; P4 vs P8, p=0.04; P4 vs P21, p=0.02). Overall, these data show that MSNs from BTBR mice display slightly earlier increases in sEPSC frequency and distinct age-dependent changes in sEPSC amplitude, but broadly equivalent timeline of maturation of sEPSC kinetics.

**Figure 1.**
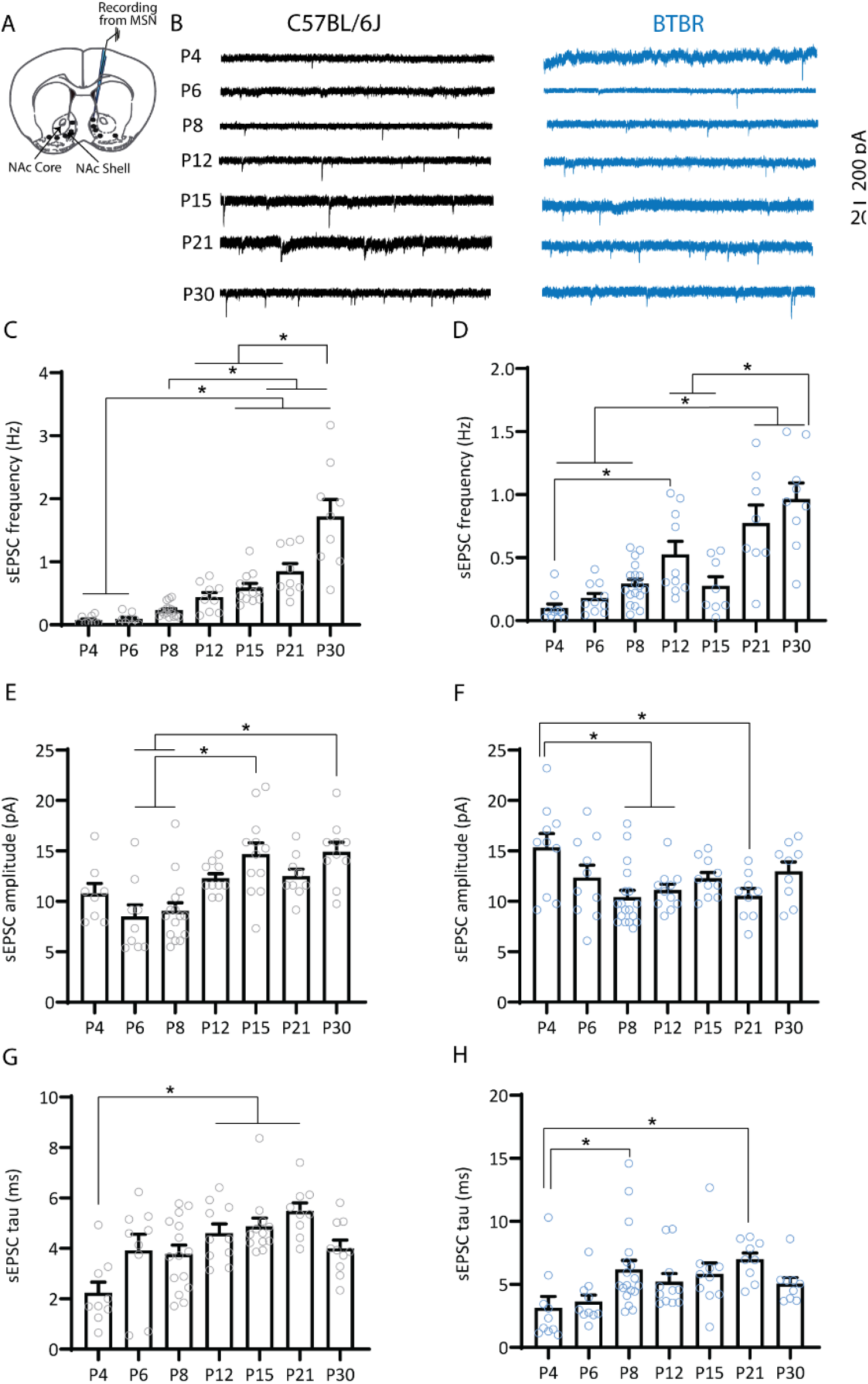
Maturation of excitatory synaptic excitation onto medium spiny neurons of the nucleus accumbens shell of C57BL/6J and BTBR mice. **A**. Schematic representation of the whole-cell patch-clamp recording protocol from an MSN of the NAc shell. **B**. Representative sEPSC traces obtained from MSNs at different postnatal ages from C57BL/6J mice (black) or BTBR mice (blue). **C-D**. Maturation of sEPSC frequency. C57BL/6J (**C**) and BTBR (**D**) mice show a progressive increase in sEPSC frequency across the first postnatal month. **E-F**. Maturation of sEPSC amplitude. C57BL/6J mice (**E**) show an age-dependent increase in sEPSC amplitude, whereas sEPSC amplitude decreases between P4 and P8-12 and P21 in BTBR mice (**F**). **G-H**. Maturation of sEPSC tau. Both C57BL6/J (**G**) and BTBR mice (**H**) show age-dependent increases in sEPSC tau. * p<0.05. C57BL/6J: P4 (9 cells/4 mice), P6 (9 cells/3 mice), P8 (15 cells/4 mice), P12 (9 cells/ 3 mice), P15 (11 cells/ 4 mice), P21 (9 cells/ 3 mice), P30 (9 cells/ 3 mice); BTBR: P4 (10 cells/3 mice), P6 (10 cells/3 mice), P8 (18 cells/5 mice), P12 (10 cells/ 3 mice), P15 (8 cells/ 3 mice), P21 (8 cells/ 3 mice), P30 (9 cells/ 3 mice).

We then examined the maturation of inhibitory inputs onto MSNs in both strains. Spontaneous inhibitory postsynaptic current (sIPSC) frequency increased significantly during the second postnatal week for both strains and plateaued from P15 (Fig. 2A-B, C57BL/6J: one way ANOVA, F_(6, 61)_ = 10.07 p<0.0001, Tukey’s multiple comparison tests: P4 vs P15, p<0.0001; P4 vs P21, p=<0.0001; P4 vs P30, p=0.004; P6 vs P15, p=0.0003; P6 vs P21, p=0.0003; P6 vs P30, p=0.01; P8 vs P15, p=0.0004; P8 vs P21, p=0.0005; P8 vs P30, p=0.03; Fig. 2A,C, BTBR, one way ANOVA, F_(6, 61)_ =5.73 p<0.0001, Tukey’s multiple comparison tests: P4 vs P15, p=0.0031, P4 vs P30, p=0.0053; P6 vs P15, p=0.007; P6 vs P30, p=0.011; P12 vs P15, p=0.017, P12 vs P30, p=0.025). In contrast, while sIPSC amplitude increased significantly and plateaued from P15 in C57BL/6J mice (Fig. 2D, C57BL/6J: one way ANOVA, F_(6, 61)_ = 7.931 p<0.0001; Tukey’s multiple comparison tests: P4 vs P15, p=0.0031; P6 vs P15, p<0.0001; P6 vs P21, p=0.0002; P8 vs P15, p=0.0009; P8 vs P21, p=0.033; P12 vs P15, p=0.041), BTBR mice only showed a slight increase at P21 (Fig. 2E, BTBR: one way ANOVA F_(6, 62)_ = 4.04, p=0.0018; Tukey’s multiple comparison tests: P8 vs P21, p=0.023; P15 vs P21, p=0.044). Interestingly, sIPSC time constant displayed a distinct maturation profile between MSNs from C57BL/6J and BTBR mice. While in C57BL/6J mice sIPSC time constant was stable across all ages tested (Fig. 2F; one way ANOVA, F_(6, 60)_ = 2.186 p=0.0567), sIPSC tau showed a significant increase between P4 and P30 in MSNs from BTBR mice (Fig. 2G; one way ANOVA, F_(6, 65)_ = 3.481 p=0.0048, Tukey’s multiple comparison tests: P4 vs P30, p=0.01). Overall, these data show broadly synchronous developmental changes in sIPSC frequency between strains, with BTBR mice showing slightly delayed increases in sIPSC amplitude and distinct maturation of sIPSC decay time.

**Figure 2.**
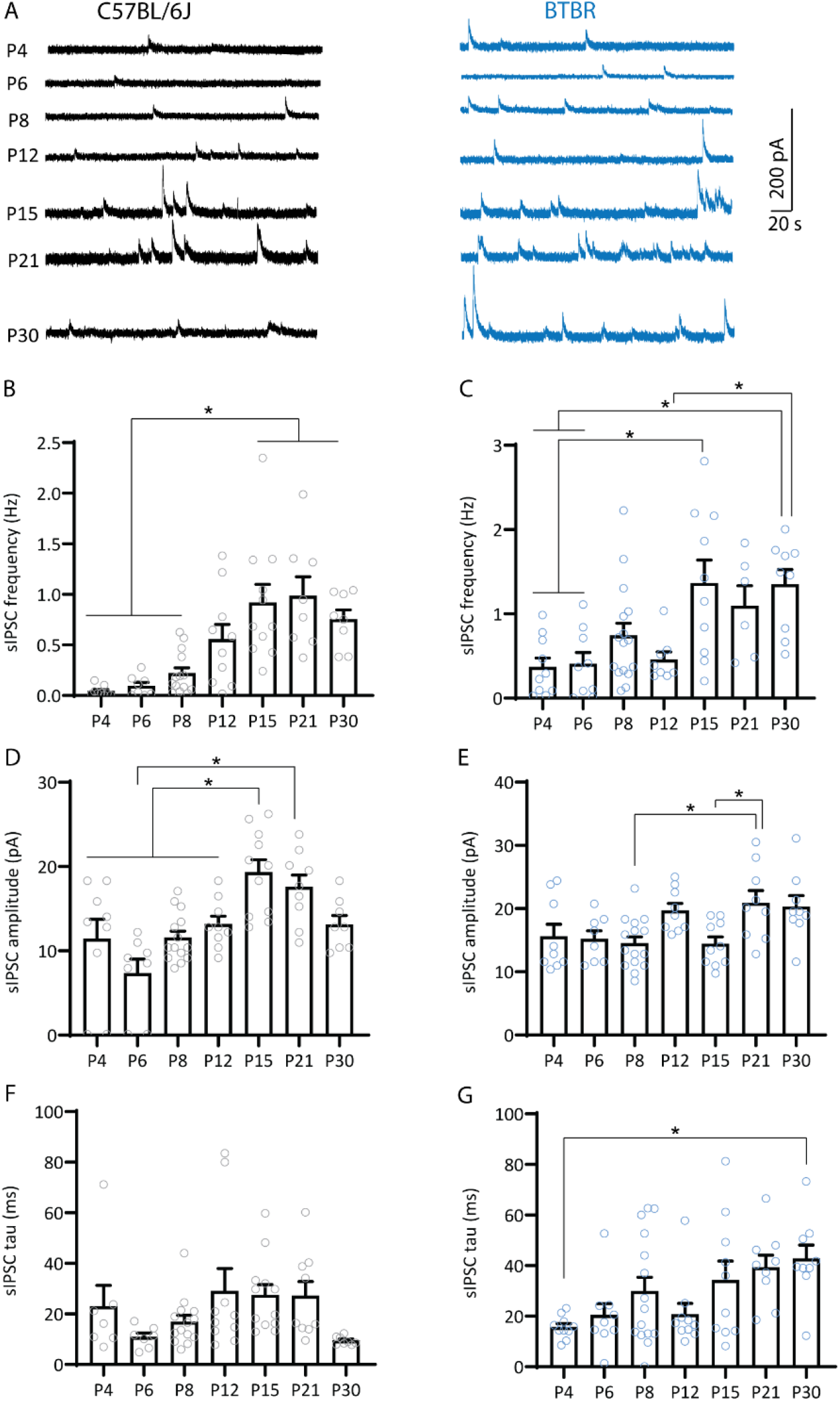
Maturation of inhibitory synaptic excitation onto medium spiny neurons of the nucleus accumbens shell of C57BL/6J and BTBR mice. **A.** Representative traces of sIPSCs obtained from MSNs in the NAc shell of C57BL/6J (black) and BTBR mice (blue). **B-C**. Maturation of sIPSC frequency. Both C57BL/6J (**B**) and BTBR mice (**C**) showed an increase in sIPSC frequency across the first postnatal month. **D-E**. Maturation of sIPSC amplitude. C57BL/6J mice (**D**) showed a slightly earlier (P15) increase in sEPSC amplitude compared to BTBR mice (**E**; P21). **F-G**. Maturation of sIPSC tau. C57BL/6J mice (**F**) showed stable sIPSC tau from P4, whereas BTBR mice (**G**) showed an increase in tau between P4 and P30. *p<0.05. C57BL/6J: P4 (9 cells/4 mice), P6 (8 cells/3 mice), P8 (14 cells/4 mice), P12 (10 cells/ 3 mice), P15 (11 cells/ 4 mice), P21 (8 cells/ 3 mice), P30 (8 cells/ 3 mice); BTBR: P4 (10 cells/3 mice), P6 (9 cells/3 mice), P8 (16 cells/5 mice), P12 (8 cells/ 3 mice), P15 (10 cells/ 3 mice), P21 (6 cells/ 3 mice), P30 (9 cells/ 3 mice).

To better understand strain differences in NAc shell MSN spontaneous excitatory and inhibitory transmission in early life, we directly compared MSNs sEPSC and sIPSC parameters between C57BL/6J and BTBR mice across ages (Fig. 3). We found that BTBR mice exhibited reduced sEPSC frequency at P30 (Fig. 3A; two way ANOVA, significant effects of age F_(6, 130)_ = 39.16, p<0.0001, strain F_(1, 130)_=5.65, p=0.02, and age x strain interaction F_(6, 130)_=4.9, p=0.0002; Sidak’s multiple comparisons P30, p<0.0001) but increased sEPSC amplitude at P4 and P6 relative to C57BL/6J mice (Fig. 3B; two way ANOVA, significant effects of age F_(1, 136)_=6.27, p<0.0001, and age x strain interaction F_(6, 136)_=4.36, p=0.0005; Sidak’s multiple comparisons P4, p=0.01; P6, p=0.04). Comparison of spontaneous inhibitory synaptic transmission revealed that BTBR mice showed higher sIPSC frequency at P8 and sIPSCs amplitude at P6, P12 and P30 compared to C57BL/6J mice (Fig. 3C-D; sIPSC frequency: two way ANOVA, significant effects of age F_(6,122)_=12.90, p<0.0001, and strain F_(6, 122)_=14.2, p=0.0003; Sidak’s multiple comparisons P8, p=0.02; sIPSC amplitude: two way ANOVA, significant effects of age F_(6, 123)_=7.08, p<0.0001, strain F_(6, 123)_=4.55, p<0.0001, and age x strain interaction F_(6, 123)_=4.55, p=0.0003; Sidak’s multiple comparisons P6, p=0.004, P12, p=0.01, P30, p=0.009). Consistent with this, comparison of within-cell differences between the frequency of spontaneous excitation and inhibition showed a shift toward synaptic inhibition at P30 in BTBR mice relative to age-matched C57BL/6J mice (Fig. 3E; two way ANOVA, significant effects of age F_(6, 120)_=5.51, p<0.0001, strain F_(6, 120)_=23.47, p<0.0001, and age x strain interaction F_(6, 120)_=3.74, p=0.0019; Sidak’s multiple comparisons P30, p<0.0001). In conclusion, these data show that MSNs from BTBR mice display higher excitatory inputs during the first postnatal week, which is reduced by P30. Inhibitory synaptic inputs onto MSNs from BTBR mice were stronger during the first, second and fourth postnatal weeks than onto MSNs from C57BL/6J mice, suggesting differences in the maturation of excitatory and inhibitory transmission in MSNs of BTBR mice. At P30, C57BL/6J and BTBR mice show opposite profiles of excitation-inhibition balance, with predominant excitation over inhibition for C57BL/6J and inhibition over excitation for BTBR mice. The temporal profile of changes in NAc synaptic transmission in BTBR mice coincides with the onset of social deficits in this strain^80^, suggesting these electrophysiological differences might contribute to the early changes in behavior.

**Figure 3.**
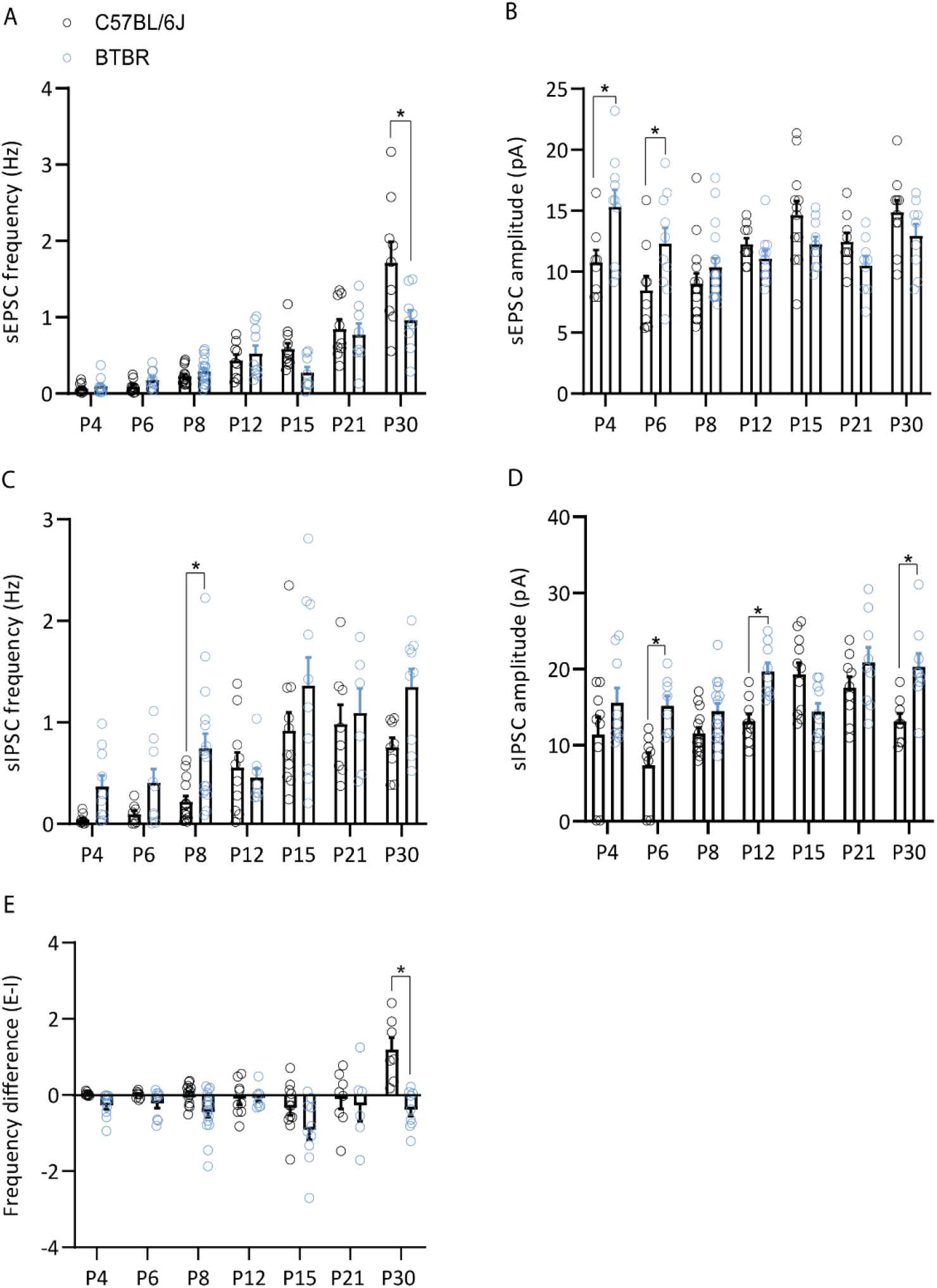
Comparison of the developmental profile of spontaneous excitatory and inhibitory synaptic transmission between C57BL/6J and BTBR mice. **A-D.** Direct comparison of age-dependent changes in NAc MSN sEPSC and sIPSC frequency and amplitude depicted in Figures 1 and 2 between C57BL/6J (black) and BTBR (blue) strains. Compared to C57BL/6J mice, BTBR mice showed reduced sEPSC frequency at P30 (**A**), increased sEPSC amplitude at P4 and P6 (**B**), increased sIPSC frequency at P8 (**C**) and increased sIPSC amplitude at P6, P12 and P30 (**D**). **E**. Within-cell comparison of the difference in the frequency of spontaneous excitation and inhibition (E-I frequency) showed a shift toward increased inhibition in P30 BTBR mice relative to C57BL6/J mice. *p<0.05. sEPSC: C57BL/6J: P4 (9 cells/4 mice), P6 (9 cells/3 mice), P8 (15 cells/4 mice), P12 (9 cells/ 3 mice), P15 (11 cells/ 4 mice), P21 (9 cells/ 3 mice), P30 (9 cells/ 3 mice); BTBR: P4 (10 cells/3 mice), P6 (10 cells/3 mice), P8 (18 cells/5 mice), P12 (10 cells/ 3 mice), P15 (8 cells/ 3 mice), P21 (8 cells/ 3 mice), P30 (9 cells/ 3 mice).

Evidence suggests specialization in the contribution of NAc input and output connections to social processing^20,91–93^, with optogenetic manipulation of prefrontal cortex (PFC)-NAc projections in particular modulating different aspects of social behavior^23,27,30,94^. To test whether PFC-NAc transmission might be altered in BTBR mice, we injected AAV-ChR2 virus into the infralimbic cortex (IL) subdivision of the PFC of C57BL/6J and BTBR mice one week prior to conducting patch clamp recordings from NAc shell MSNs at P15 and P30 (Fig. 4A). IL-expressing ChR2 terminals onto MSNs were stimulated using a brief light-pulse (5ms) through the objective, and glutamate release probability at IL-NAc synapses was examined by measuring paired-pulse ratios (PPR). Light-evoked EPSCs from MSNs of BTBR mice showed higher PPR at both P15 and P30 compared to C57BL/6J mice, indicating reduced presynaptic glutamate release probability at IL-NAc synapses (Fig. 4B-C; unpaired t tests, P15: t=2.13, p=0.04; P30: t=5.91, p<0.0001).

**Figure 4.**
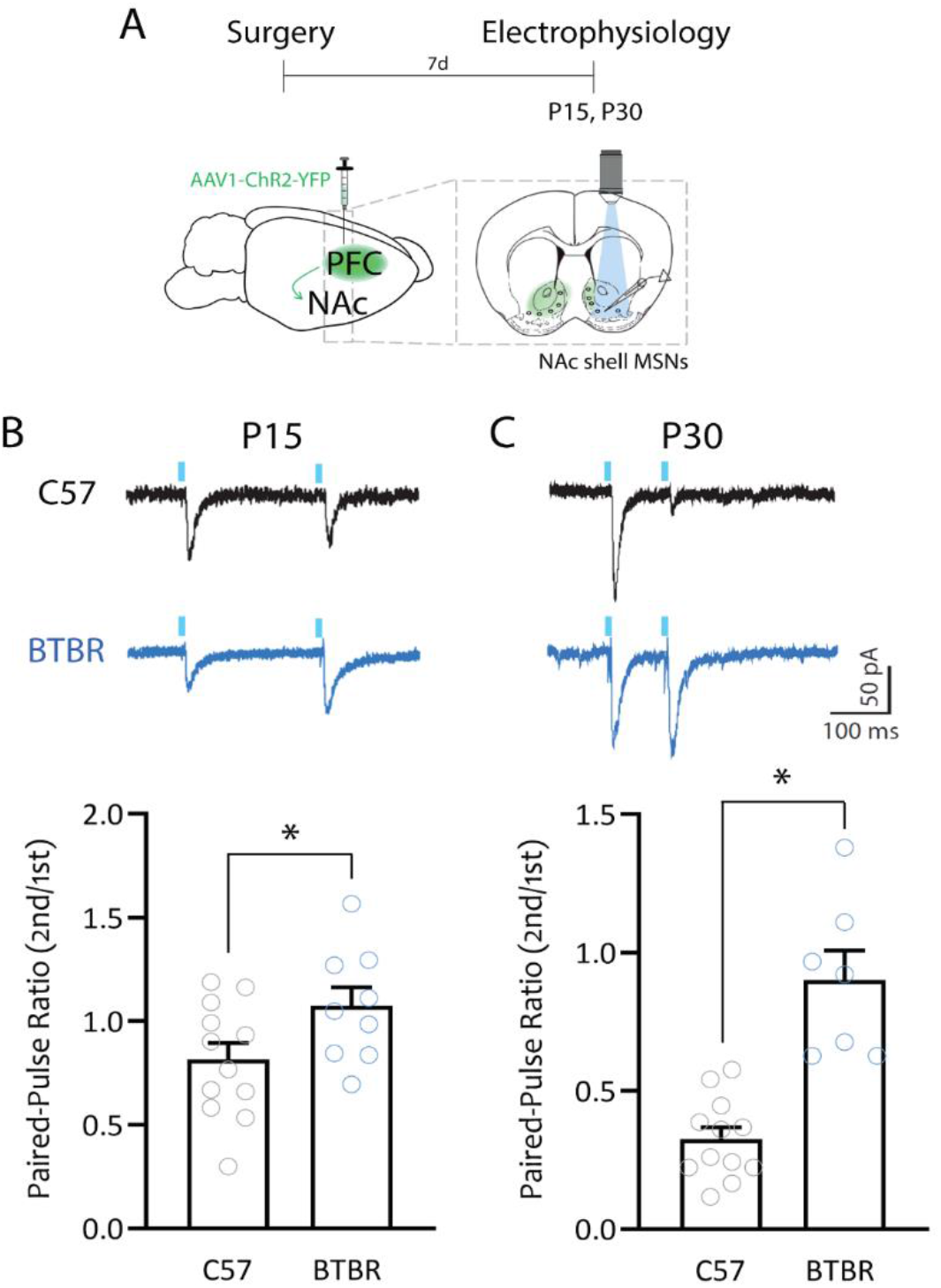
BTBR mice show decreased PFC-NAc glutamate release probability in early life. **A.** Schematic of experimental design. **B.** Top: P15, representative traces of light-evoked EPSC during paired-pulse stimulation (250 ms) from MSNs of C57BL/6J mice (black) and BTBR mice (blue). Bottom: summary plot showing increased PPR in MSNs from BTBR mice compared to MSNs from C57BL/6J mice. **C**. Top: P30, representative traces of light-evoked EPSC during paired-pulse stimulation (100 ms) from MSNs of C57BL/6J mice (black) and BTBR mice (blue). Bottom right: summary graph showing increased PPR in MSNs from BTBR mice compared to those from C57BL/6J mice. * p<0.05. C57BL/6J: P15 (12 cells/ 6 mice), P30 (12 cells/ 4 mice); BTBR: P15 (9 cells/ 4 mice), P30 (7 cells/ 4 mice).

Given the temporal coincidence of alterations in NAc synaptic transmission and behavior in the BTBR strain, targeting intervention strategies to this window may improve treatment outcomes. Informed by our electrophysiological data showing changes in NAc spontaneous transmission as early as P4 (Figs 1–3), we decided to target a well-established rescue strategy for animal models of ASD, the mTORC1 antagonist rapamycin^95^, to early development, starting at P4. We first confirmed that BTBR mice displayed social interaction deficits in our lab by testing them on a three-chamber social interaction test, followed by a social memory test. We tested animals at P30, a crucial time in social development^96^. In the social interaction test, BTBR mice spent less time with the social target and more time in the empty compartment, compared to C57BL/6J mice (Fig. 5A-B; two-way RM ANOVA, significant effect of chamber F_(1.536, 90.61)_= 38.79, p<0.0001 and chamber x strain interaction F_(2,118)_= 16.71, p<0.0001; Sidak’s multiple comparisons test, empty p=0.0074, middle p=0.0030, and social p=0.0006 chambers).

**Figure 5.**
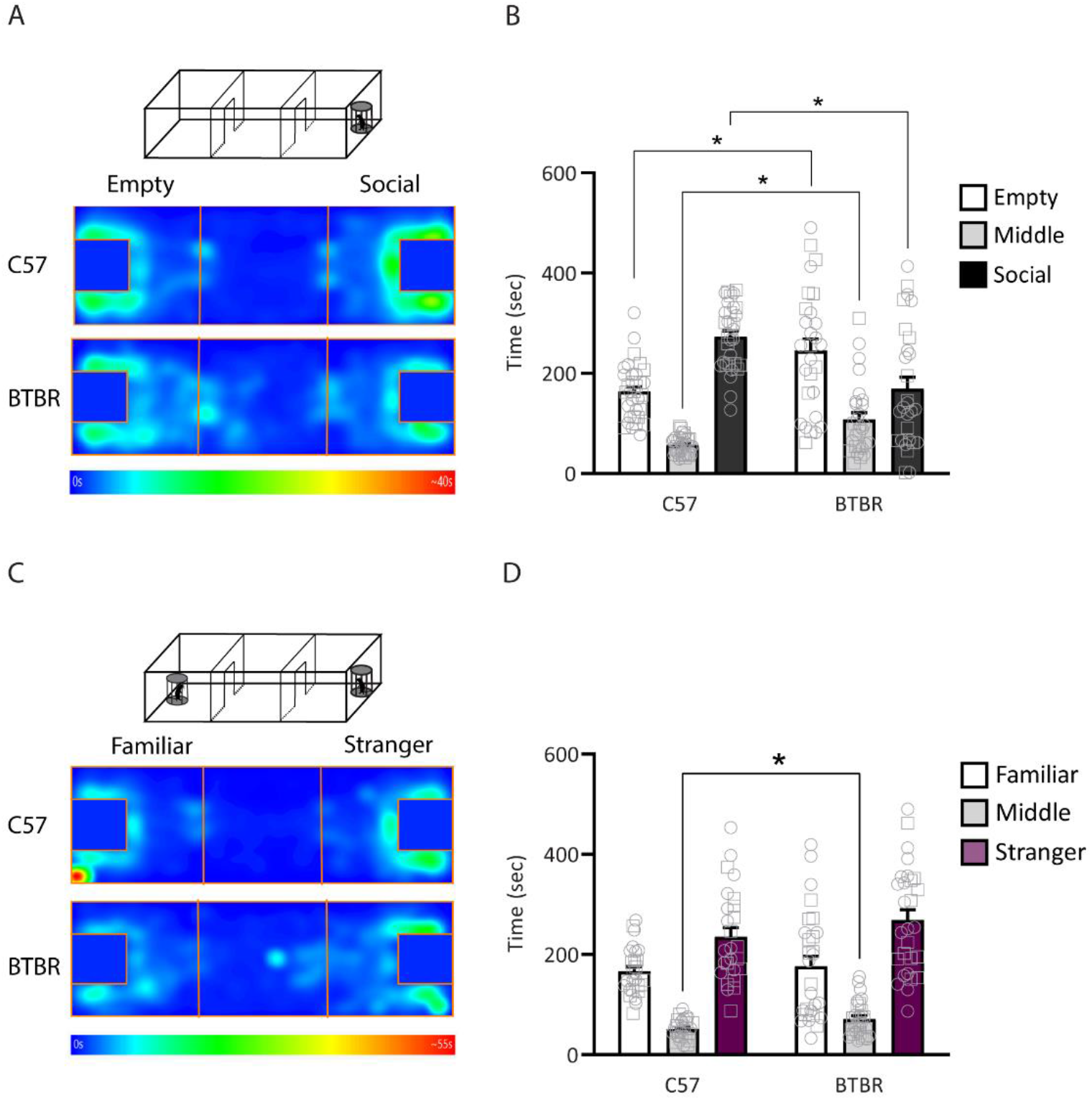
BTBR mice show social interaction deficits but no change in social memory at P30. C57BL/6J and BTBR underwent the three-chamber social interaction test followed by a social memory test at P30. **A.** Schematic of behavioral testing and heatmaps of time spent in each chamber during testing. **B.** Three chamber social interaction test. Time spent in the chamber with an empty enclosure (empty), middle chamber and chamber with an age-, strain- and sex-matched stranger mouse in the enclosure (social). C57BL/6J mice spent significantly more time exploring the social chamber (Sidak’s post hoc test, Empty vs. Middle t_32_= 10.63 p<0.0001, Empty vs. Social t_32_= 7.365 p<0.0001, Middle vs. Social t_32_= 18.48 p<0.0001), whereas BTBR mice showed no significant preference for the social chamber (Sidak’s post hoc test, Empty vs. Middle t_27_= 4.843 p=0.0001, Empty vs. Social t_27_= 1.76 p=0.2458, Middle vs. Social t_27_= 2.045 p=0.1446). C57BL/6J, n = 33 (17 females, 16 males); BTBR, n = 28 (17 females, 11 males). **C.** Social memory test. Time spent exploring the chamber with the same mouse from the previous test (familiar), middle chamber, and chamber with a novel, age, strain and sex-matched stranger mouse in the enclosure (stranger). Both C57BL/6J and BTBR mice showed a preference for the stranger mouse (Sidak’s post hoc test, C57BL/6J: Stranger vs. Middle t_25_= 9.019 p<0.0001, Stranger vs. Familiar t_25_= 3.366 p=0.0074, Middle vs. Familiar t_25_= 11.14 p<0.0001; BTBR: Stranger vs. Middle t_30_= 5.282 p<0.0001, Stranger vs. Familiar t_30_= 2.533 p=0.0494, Middle vs. Familiar t_30_= 2.237 p=0.0955. C57BL/6J, n = 26 (13 females, 13 males); BTBR, n = 31 (14 females, 17 males). No sex differences were found for either strain in the three-chamber social interaction test (p>0.64) or social memory test (p>0.11). Male individual datapoints are depicted as squares, and female datapoints as circles for transparency. *p<0.05.

While BTBR social interaction deficits consistent with ours have been widely reported^79,97^, social memory deficits in the BTBR strain have a more conflicting literature^75,98,99^. We found no differences in social memory, i.e. preference for the novel, unfamiliar mouse, between C57BL/6J and BTBR mice (Fig. 5C-D; two-way RM ANOVA, significant effect of chamber F_(2, 110)_= 39.63, p<0.0001 and strain F_(1,55)_= 10.23 p=0.0023; Sidak’s multiple comparisons test showed significant differences between strains only in the middle chamber t=2.775 p=0.0268; p>0.84 for other chambers). Importantly, although our behavioral data showed no sex differences in social interaction (p>0.64) or social memory (p>0.11), replicating a widely reported absence of sex differences in social behavior in these strains^79,84,85,89^, we presently cannot exclude the possibility of sex differences in the underlying maturation of NAc synaptic transmission.

To test whether the increased synaptic excitation and inhibition onto MSNs from BTBR mice starting at the first postnatal week could signal the start of a sensitive period in the BTBR strain, we targeted rapamycin treatment, a drug known to rescue social deficits in other mouse models of ASD^65,66^, from P4 to P8, and tested animals for social preference at P30 (Fig. 6A). Rapamycin treatment led to a significant increase in time spent in the social chamber (Fig. 6B; two-way RM ANOVA, significant effect of chamber F_(2, 48)_= 22.42, p<0.0001, and chamber x treatment interaction F_(2,48)_= 9.436, p<0.0003; Sidak’s multiple comparisons test show significant differences between vehicle and rapamycin treatment in the empty p=0.0293 and social p=0.0002 chambers), indicating a rescue of social preference in BTBR mice treated with rapamycin from P4-P8. We saw no difference in distance travelled between vehicle- and rapamycin-treated groups (Fig. 6C; unpaired t test, t_19_=0.2272, p=0.8227), suggesting that the social interaction effects did not stem from changes in motor activity. These data indicate that early rapamycin treatment was able to rescue BTBR social deficits at adolescence. As an additional validation, we confirmed that early rapamycin treatment decreased phosphorylation of S6 ribosomal protein, a commonly used measure of mTOR pathway activity^100^, in P30 BTBR mice (Supplementary Fig. 1A-B; unpaired t test, t_6_=3.2, p=0.01).

**Figure 6.**
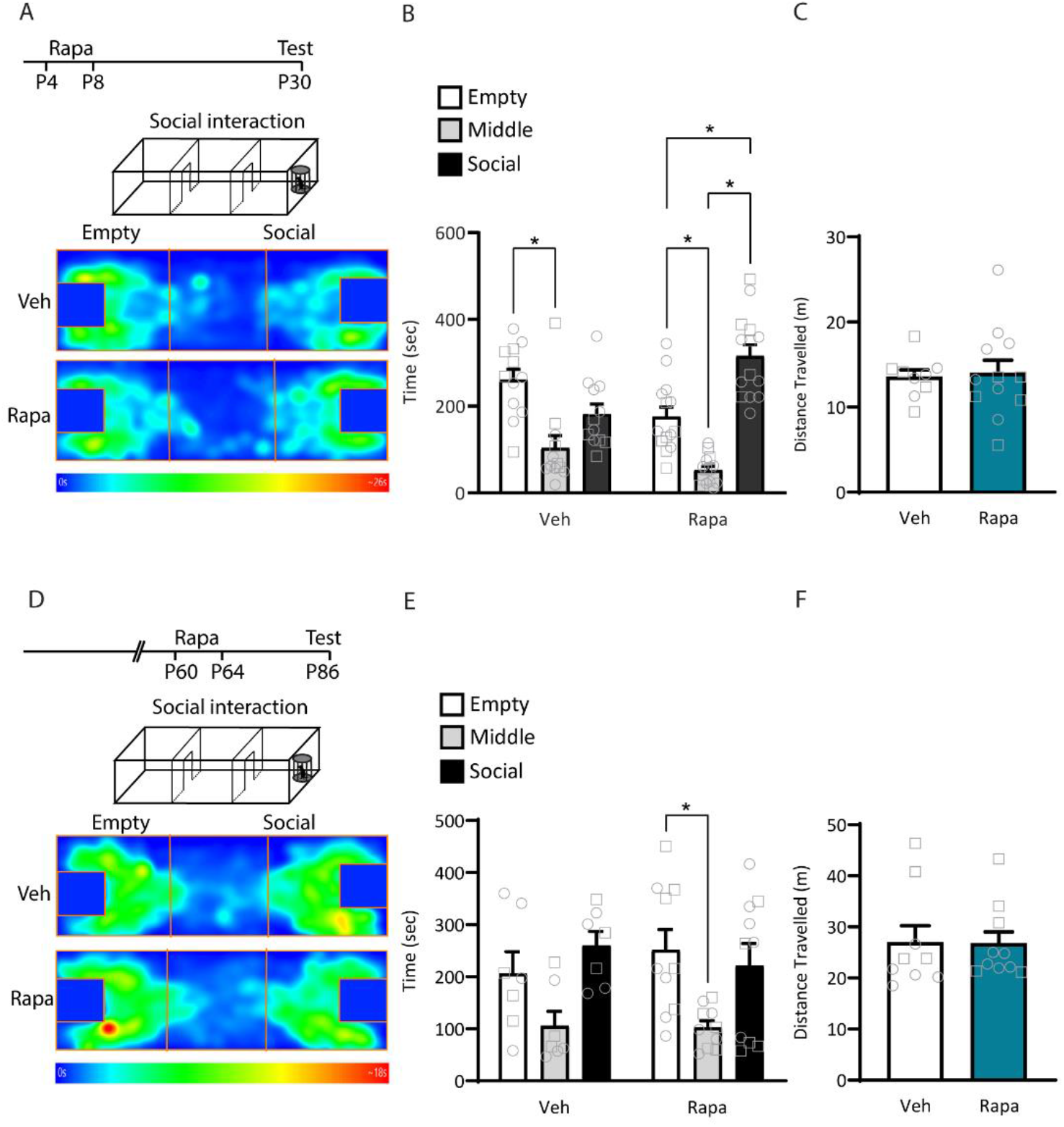
Rapamycin treatment in infancy, but not adulthood, rescues BTBR mice social interaction deficits. **A-C.** BTBR mice were treated with rapamycin or vehicle from P4-P8 and tested in the three-chamber social interaction test at P30. **A.** Schematic of experimental design and heatmaps of time spent in each chamber during testing. **B.** Three chamber social interaction test. Time spent in the chamber with an empty enclosure (empty), middle chamber and chamber with an age-, strain- and sex-matched stranger mouse in the enclosure (social). Rapamycin-treated mice showed a preference for the chamber with the social target (Sidak’s post hoc test, Empty vs. Middle t_48_= 3.334 p=0.005, Empty vs. Social t_48_= 3.799 p=0.0012, Middle vs. Social t_48_= 7.132 p<0.0001), which is absent in vehicle-treated BTBR mice (Sidak’s post hoc test, Empty vs. Middle t_48_= 3.937 p=0.0008, Empty vs. Social t_48_= 1.982 p=0.1513, Middle vs. Social t_48_= 1.954 p=0.1601). **C.** Distance travelled in the social interaction test. There was no difference between vehicle- and rapamycin-treated mice. Vehicle-treated BTBR (VEH), n = 12 (7 females, 5 males); Rapamycin-treated BTBR (RAPA), n = 14 (8 females, 6 males). No sex differences were found for either group (p>0.516). Male individual datapoints are depicted as squares, and female datapoints as circles for transparency. **D-F**. BTBR mice were treated with rapamycin or vehicle from P60-P64 and tested in the three-chamber social interaction test at P86. **D.** Schematic of experimental design and heatmaps of time spent in each chamber during testing. **E.** Three chamber social interaction test. Time spent in the chamber with an empty enclosure (empty), middle chamber and chamber with an age, strain and sex-matched stranger mouse in the enclosure (social). Neither vehicle or rapamycin treated mice showed a preference for the chamber with the social target over the empty chamber (VEH: Sidak’s post hoc test, Empty vs. Middle t_34_= 2.214 p=0.097, Empty vs. Social t_34_= 2.012 p=0.148, Middle vs. Social t_34_= 4.22 p=0.0005; RAPA: Empty vs. Middle t_34_= 2.72 p=0.030, Empty vs. Social t_34_= 0.557 p=0.93, Middle vs. Social t_34_= 2.16 p=0.11). **F.** Distance travelled in the social interaction test. There was no difference between vehicle- and rapamycin-treated mice. Vehicle-treated BTBR (VEH), n = 9 (5 females, 4 males); Rapamycin-treated BTBR (RAPA), n = 10 (5 females, 5 males). No sex differences were found for either group (p>0.1185). Male individual datapoints are depicted as squares, and female datapoints as circles for transparency. *p<0.05.

A study by Burket and colleagues had shown that a four-day rapamycin treatment starting at P28 also rescued social deficits in BTBR mice tested 1h after the last injection^101^. To test whether the efficacy of the rapamycin rescue was tied to its timing during early life, we injected an additional cohort of BTBR mice with the same rapamycin regimen and interval between treatment and testing, but starting at P60 (Fig. 6D). We found that rapamycin treatment starting at early adulthood did not affect the time spent in the social chamber (Fig. 6E; two-way RM ANOVA, significant effect of chamber F_(2, 34)_= 11.32, p=0.0002; Sidak’s multiple comparisons test show no significant differences between vehicle and rapamycin treatment in any of the chambers) or distance travelled (Fig. 6F; unpaired t test, t_17_=0.052, p=0.9591). Overall, these data indicate that P60 rapamycin treatment did not rescue social interaction deficits in BTBR mice, suggesting that early intervention with rapamycin is necessary for improving social interaction in the BTBR strain.

## Discussion

In this study, we compared the maturation of NAc shell MSN spontaneous synaptic transmission in C57BL/6J and BTBR strains from infancy to adolescence. In both strains, MSNs displayed excitatory and inhibitory spontaneous currents by P4, suggesting the presence of functional excitatory and inhibitory synapses in the NAc early during development. Excitatory and inhibitory inputs onto MSNs developed gradually up to P12 and underwent a marked maturation by P15, when levels plateaued. These data suggest that synaptic inputs onto NAc MSNs establish early during development with a slow maturation during the first 10-12 postnatal days, followed by a potentiation of synaptic inputs onto MSNs by P15. This is consistent with other reports showing stable NAc paired pulse ratio (PPR) of AMPA-mediated EPSCs from juvenility^50^, and changes in AMPAR- and NMDAR-mediated EPSCs up to the second postnatal week (but decrease of NMDA-EPSC later during development)^52^.

Overall, we found age-dependent changes in the maturation of spontaneous excitatory and inhibitory transmission within and between strains, with BTBR mice showing increased excitation during the first postnatal week, which decreased by P30, as well as increased spontaneous inhibition across early development. Additionally, BTBR mice display IL-NAc paired-pulse facilitation at P15 and P30 compared to C57BL/6J mice, suggesting reduced presynaptic glutamate release from IL terminals onto MSNs in BTBR mice. The difference in PPR between C57BL/6J and BTBR mice at P15 and P30 was observed using long (250 ms) and short (100 ms) interstimulus intervals, respectively, which might be influenced by developmental differences in presynaptic short-term plasticity, as reported previously^102–104^

In the *Shank3B^-/-^* mouse model of ASD, an accelerated maturation of striatal spiny projecting neurons (SPNs) of the dorsomedial striatum (DMS) was observed during early developmental stages (third and fourth postnatal weeks), followed by a decrease in corticostriatal activity during adulthood (P60)^105,106^ - a deficit that was attributed to an overactivation of cortical circuits. Moreover, the authors found that increasing anterior cingulate cortex (ACC) synaptic drive to SPNs of DMS at P8 significantly increased SPN mEPSC frequency, whereas increasing ACC activity from P15 and older reduced SPN mEPSC frequency^106^, suggesting a developmental switch in the effect of cortical hyperactivity on SPN synaptic transmission. While our study showed that MSNs from BTBR mice receive weaker synaptic inputs from IL at P15 and P30 relative to MSNs from C57BL/6J mice, it remains unknown whether a similar switch occurs in their response to IL hypofunction.

Given the developmental timeline of NAc shell MSN spontaneous transmission, we hypothesized that early postnatal development might constitute a critical period for the efficacy of rescue manipulations correlating with improved outcomes. To test this, we subjected BTBR mice to rapamycin treatment in infancy (P4-P8) or adulthood (P60-64) and tested its effect on social preference in a three-chamber social interaction test. We found that rapamycin treatment in infancy, but not adulthood, reversed the social interaction deficits in BTBR mice. Our data point to a critical period for the efficacy of rapamycin treatment in BTBR mice from P4-P8 which closes by P60. Work by Burket and colleagues showed that rapamycin rescues BTBR social interaction deficits when administered between P28-P31, with animals tested one hour after the last injection^101^. While this study did not probe the long-term effects of rapamycin treatment on social behavior, it reinforces the value of rapamycin as an effective rescue intervention in BTBR mice. Furthermore, it points to a longer window of efficacy for rapamycin treatment in this strain, likely spanning from infancy to early adolescence. Consistent with this, we found differences in NAc spontaneous transmission between BTBR and C57BL/6J mice up to P30. In the rat, changes in NAc intrinsic excitability and excitatory transmission predominantly occur within the first three postnatal weeks, reaching adult levels by P35 or earlier^50,51^. A study by Zhang and Warren saw changes in rat NAc excitatory transmission stabilizing by P15, with the amplitude of NMDA responses peaking from P9-12, which they propose is a critical period for this brain region^52^. Additionally, Liu and colleagues reported higher sEPSC frequency and amplitude, as well as lower halfwidth, rise and decay time in the NAc shell of P21-P28 rats compared to adults^107^, further pointing to the first postnatal month as a developmental window of heightened changes in NAc synaptic transmission.

Nardou and colleagues had described a critical period for social reward learning in the NAc^108^, which peaks at P42, and closes by P98^108^. This learning window coincides with age-dependent changes in oxytocin-induced long-term synaptic depression in the NAc^16,108^. Interestingly, NAc oxytocin signaling also modulates social approach^21^, suggesting its signaling and developmental time course might also influence social interaction. Another NAc microcircuit regulator, μ-opioid receptor (MOR), modulates social interaction and NAc synaptic transmission^109^, as well as social behavior in juveniles such as play behavior^110,111^ and social preference^112^. Furthermore, mice lacking a functional MOR gene (Oprm1^-/-^mice), an ASD mouse model, display increased oxytocin receptor expression in the NAc, with their social deficits reversed by oxytocin treatment^113^. NAc MOR levels^114,115^ and function^116^ also change across development, suggesting that the interplay between NAc MOR and oxytocin might contribute to the shaping of social behavior in early life.

While a wide array of acute or short-term interventions rescue social interaction deficits in adult ^75,97,125–134,117,135–140,118–124^ and young^141–146^ BTBR mice, very few probed their long-term efficacy. Pharmacological rescue interventions yielding long-term rescue of BTBR social deficits predominantly target mice up to the third postnatal week^147–151^, with effects lasting until adulthood. To our knowledge, no other study compared the efficacy of a rescue intervention across different ages in BTBR mice. In an interesting study^41^, deletion of Shank3 in the NAc at a similar time period to ours (<P6) triggered social deficits in adulthood that were not replicated when the manipulation started at P90, corroborating the existence of a critical period for NAc-associated behavior and susceptibility to long-lasting interventions. Notably, the NAc is not the only region contributing to the regulation of social behavior in the BTBR strain, as seen with adult manipulations of cerebellum^152^ and PFC^153^, as well as changes in spontaneous transmission in hippocampus^140^. While we saw decreased presynaptic efficacy in the PFC-NAc pathway of BTBR mice, its precise contribution to behavioral deficits in this strain remains unclear.

Our data delineate the maturation of spontaneous synaptic transmission in the NAc shell across early postnatal development, uncovering distinct early maturation signatures between C57BL/6J and BTBR mice, strains that display distinct patterns of NAc-mediated social behavior. These synaptic changes coincide with key developmental changes in social communication, play, affiliative and social approach behavior^154–158^, and abridge a period in which environmental conditions and stress bear a disproportionate impact in shaping future social behavior^159–163^. Accordingly, we revealed a restricted time window for the rescue of social interaction deficits in BTBR mice by rapamycin treatment. These findings emphasize the importance of matching experimental conditions to clinically relevant timelines, and highlight the power of leveraging research in early development toward precisely targeted interventions that maximize positive outcomes.

## Acknowledgements

We thank Mehreen Inayat and Ryan Appings for help with mouse breeding and Jennifer Wilkin for help with the design of figure schematics. We acknowledge resources and support from the Centre for the Neurobiology of Stress (CNS) Core Facility at the University of Toronto Scarborough.

