## Supplementary Figure 1 for "Maturation of Nucleus Accumbens Synaptic Transmission Signals a Critical Period for the Rescue of Social Deficits in a Mouse Model of Autism Spectrum Disorder"

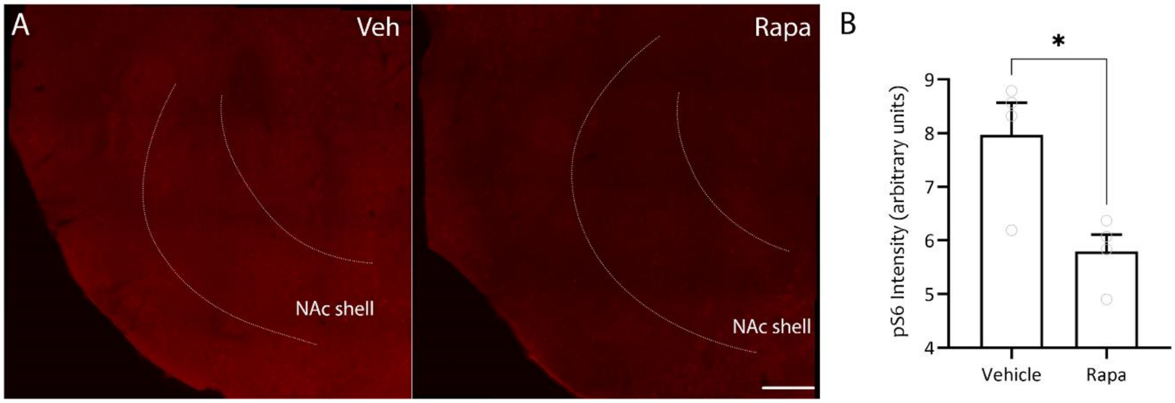


**Supplementary Figure 1. Rapamycin treatment decreases phosphoS6 levels in the NAc of BTBR mice at P30.** BTBR mice were treated with rapamycin or vehicle from P4-P8, kept in their homecage and then perfused at P30 for brain processing and immunohistochemistry against phosphoS6. **A**. Representative images of phosphoS6 staining in the NAc shell. **B**. Quantification of phosphoS6 intensity shows that rapamycin treatment reduced phosphoS6 levels at P30. Scale bar = 250μm, n=4 vehicle, 4 BTBR. *$p<0.05$.
